# Transcriptional response of signalling pathways to SARS-CoV-2 infection in normal human bronchial epithelial cells

**DOI:** 10.1101/2020.06.20.163006

**Authors:** Ak Enes, Pınar Pir

**Affiliations:** Department of Bioengineering, Faculty of Engineering, Gebze Technical University, Kocaeli, Turkey

**Keywords:** SARS-CoV-2, H1N1, NHBE, Signaling pathways, Transcriptomics

## Abstract

SARS-CoV-2 virus, the pathogen that causes Covid-19 disease, emerged in Wuhan region in China in 2019, infected more than 4M people and is responsible for death of at least 300K patients globally as of May 2020. Identification of the cellular response mechanisms to viral infection by SARS-CoV-2 may shed light on progress of the disease, indicate potential drug targets, and make design of new test methods possible.

In this study, we analysed transcriptomic response of normal human bronchial epithelial cells (NHBE) to SARS-CoV-2 infection and compared the response to H1N1 infection. Comparison of transcriptome of NHBE cells 24 hours after mock-infection and SARS-CoV-2 infection demonstrated that most genes that respond to infection were upregulated (320 genes) rather than being downregulated (115 genes).While upregulated genes were enriched in signalling pathways related to virus response, downregulated genes are related to kidney development. We mapped the upregulated genes on KEGG pathways to identify the mechanisms that mediate the response. We identified canonical NFκB, TNF and IL-17 pathways to be significantly upregulated and to converge to NFκB pathway via positive feedback loops. Although virus entry protein ACE2 has low expression in NHBE cells, pathogen response pathways are strongly activated within 24 hours of infection. Our results also indicate that immune response system is activated at the early stage of the infection and orchestrated by a crosstalk of signalling pathways. Finally, we compared transcriptomic SARS-CoV-2 response to H1N1 response in NHBE cells to elucidate the virus specificity of the response and virus specific extracellular proteins expressed by NHBE cells.

## 1. INTRODUCTION

SARS-CoV-2 virus is a member of *Coronaviridae* family. This family of viruses includes large single-stranded and positive-sense RNA viruses. Infection by *Coronaviridae* family can cause gastrointestinal and hepatic disorders but the most severe effect is on respiratory system of the patients (Weston and Frieman, 2020). SARS-CoV-2, initially known as 2019-nCoV, was detected in patients in Wuhan region of China in 2019, the disease Covid-19 caused by SARS-CoV-2 infection was announced to be a pandemic by WHO in February 2020 (Yi et al., 2020).

Symptoms of SARS-CoV-2 infection varies among patients but according to clinical reports, high fever (98%) is the major symptom of Covid-19. This is followed by other symptoms such as dry cough (%76), myalgia or fatigue (44%) (Wu et al., 2020). Symptoms are significantly milder in infants and children when compared to symptoms in older people, and asymptomatic infection can be seen in younger age ranges whereas patients older than 60 years have higher risk of developing severe symptoms (Singhal, 2020). The main cause of deaths from SARS-CoV-2 is acute respiratory distress syndrome (ARDS).

Inflammation mechanisms are vital for the defence against pathogens. A crosstalk between immune system, nervous system and coagulative-fibrinolytic pathways orchestrated by “inflammatory mediators” regulate the response to inflammation. Inflammatory mediators can be classified into two types, one type is cell-derived and the other one is plasma-derived, cell-derived mediators includes cytokines (Coskun Benlidayi, 2019). SARS-CoV leads to increased production of the cytokines and chemokines in the cells against virus infection (Dosch et al., 2009) and a similar response is observed in SARS-CoV-2 infection. Synthesis of cytokines are regulated by signalling pathways such as NFκB, IL-17 which are known as proinflammatory signalling pathways (Lawrence, 2009; McGeachy et al., 2019). Excessive production of cytokines is a complication that effects the severity of the infection via hyperinflammation. Recently, integration of transcriptomic profile with metabolic networks has demonstrated that metabolic networks are also potentially dysregulated in response to SARS-CoV-2 infection (Karakurt and Pir, 2020).

Here we investigate the effect of SARS-CoV-2 infection on signalling pathways in NHBE cells to elucidate the infection mechanisms that may help identification of potential drug targets or design of new test methods. We analysed the functions of significantly changed genes via enrichment of them in GO biological process terms and KEGG signalling pathways, mapping of the genes on pathways indicated that the crosstalk between the pathways are mediated by positive feedback loops. We compared the genes that were upregulated in SARS-CoV-2 infection in NHBE cells to those that were upregulated in H1N1 infection to be able to identify the SARS-CoV-2-specific response when compared to influenza.

## 2. MATERIALS AND METHODS

Two sets of transcriptome data from normal human bronchial epithelial (NHBE) cell line was used (Accession number: GSE147507, Blanco-Melo et al., 2020) in this study. The RNA-Seq count matrix was downloaded from National Center for Biotechnology Information (NCBI) Gene Expression Omnibus (GEO) database. One set is composed of samples collected 24 hours after mock or SARS-CoV-2 virus infection (3 replicates) and other set is composed of samples collected 12 hours after mock or H1N1 virus infection (4 replicates). NHBE cells that received the mock treatment are referred to as ‘control samples’ throughout this report.

Statistical analysis of the RNA-Seq data was performed in R version 3.6.0. Principal-component analysis (PCA) was applied on normalized and standardized count matrix to inspect the structure of the data in reduced dimensions. Differential gene expression analysis was performed to compare infected and control samples using DESeq2 R package which applies Wald Test to determine significant differences between means of two groups of samples (Love et al., 2014). 0.05 was chosen as adjusted p-value threshold after applying Benjamini Hochberg correction to p-values from Wald test (Benjamini and Hochberg, 1995). Enrichment analysis of the significant changed genes in GO biological process terms and KEGG pathways were performed using Panther (Mi et al., 2018) and g:Profiler (Raudvere et al., 2019) online enrichment analysis tools respectively, Fisher’s Exact Test was applied to calculate the p-values for significant enrichment and p-values were corrected for multiple testing by using Benjamini-Hochberg method. Visualization and summarization of the gene ontology results were performed by REVIGO (Supek et al., 2011) tool. To map the significantly changed genes onto signalling pathways, KEGG Pathway Database GUI (Qiu, 2013) was used.

## 3. RESULTS

Prior to downstream transcriptome data analysis, PCA was done on the dataset which is composed of 3 control / 3 SARS-CoV-2 infected and 4 control / 4 H1N1 infected samples and scores on first two principal components were plotted (Figure 1). Variance explained by PC1 is 80% and variance explained by PC2 is 9%, PC1 separates the two batches, whereas control samples and infected samples are separated on PC2. Hence, main source of variation in the dataset, apart from the batch effect, is infection by either SARS-CoV-2 or H1N1. Next, differential gene expression analysis was applied to identify the upregulated and downregulated genes in response to by SARS-CoV-2 infection. 435 genes were identified as significantly changed (adjusted p-value < 0.05), 320 genes were upregulated and 115 genes were downregulated. These two groups of genes were further investigated to identify the biological processes and pathways that significantly responded to virus infection.

**Figure 1.**
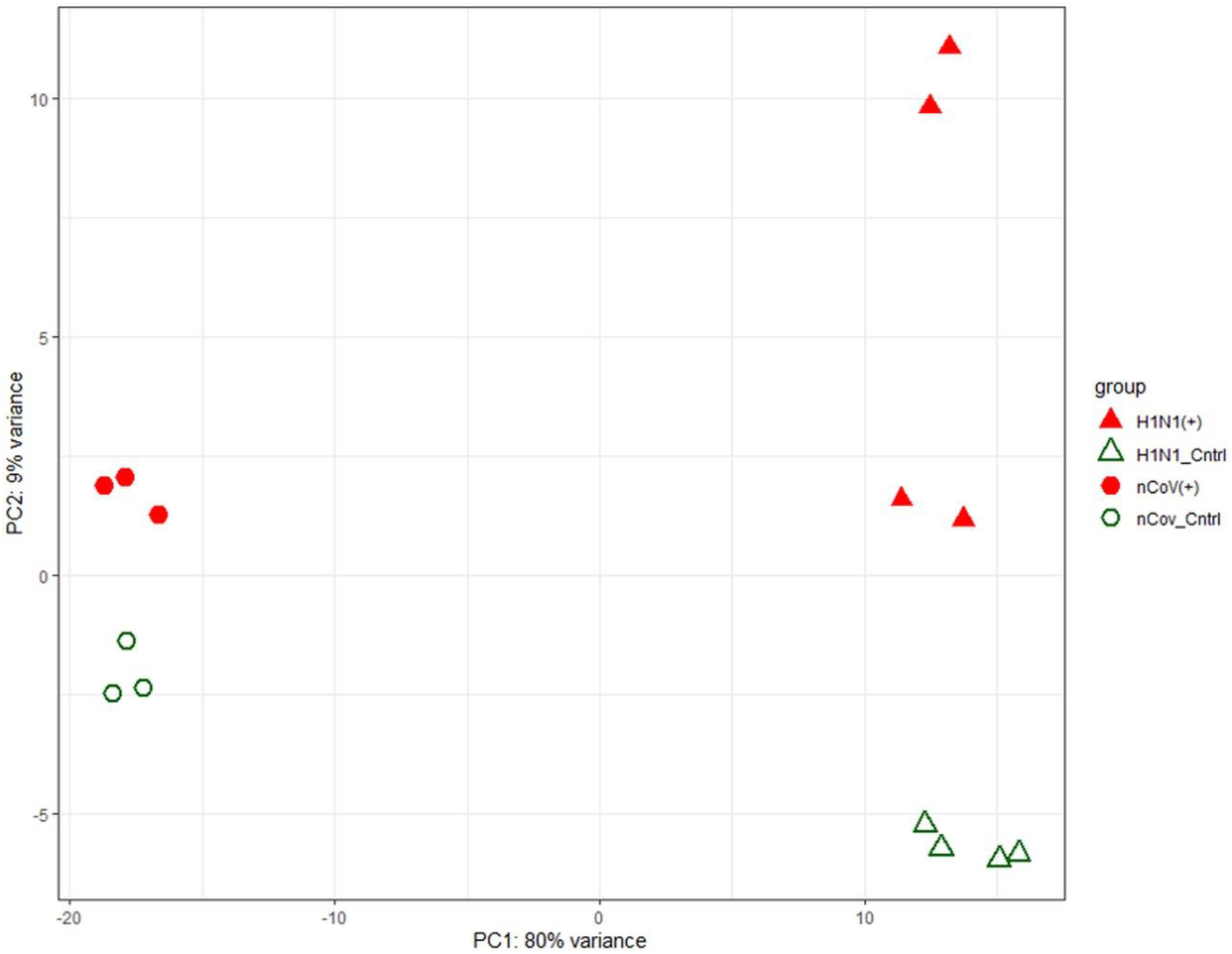
PCA of gene expression in control (mock infected), SARS-CoV-2 infected and H1N1 infected samples.

GO terms enriched with upregulated genes were related with immune system and response the external stimulus (Table 1 and Supplementary Figure 1). Most significant three GO terms were immune system process (Padj = 1.4E-36), defence response (Padj = 3.1E-36), response the cytokine (Padj = 8.9E-36). Significantly enriched GO terms were not specific enough to identify the mechanism of the cellular response, whereas KEGG pathway enrichment analysis (Table 2) allowed us to elucidate the specific pathways that responded to infection. To better understand the transcriptional changes in signalling pathways in response to SARS-CoV-2 infection, upregulated and downregulated genes were mapped onto pathways with most significant response, namely, IL-17 signalling pathway (Padj = 1.3E-17, Figure 2), NF-kappa B signalling pathway (NFκB) (Padj = 1.3E-14, Figure 3), Influenza A signalling pathway (Padj = 2.2E-14, Figure 4), TNF signalling pathway (Padj = 9.4E-14, Figure 5) and rheumatoid arthritis (Padj = 1.3E-09, Supplementary Figure 2) using KEGG Pathway Database GUI. Of note, none of the members or targets of those pathways were significantly downregulated in NHBE cells in response to SARS-CoV-2 infection, except for BLNK, B cell linker protein, in NFκB pathway.

**Figure 2.**
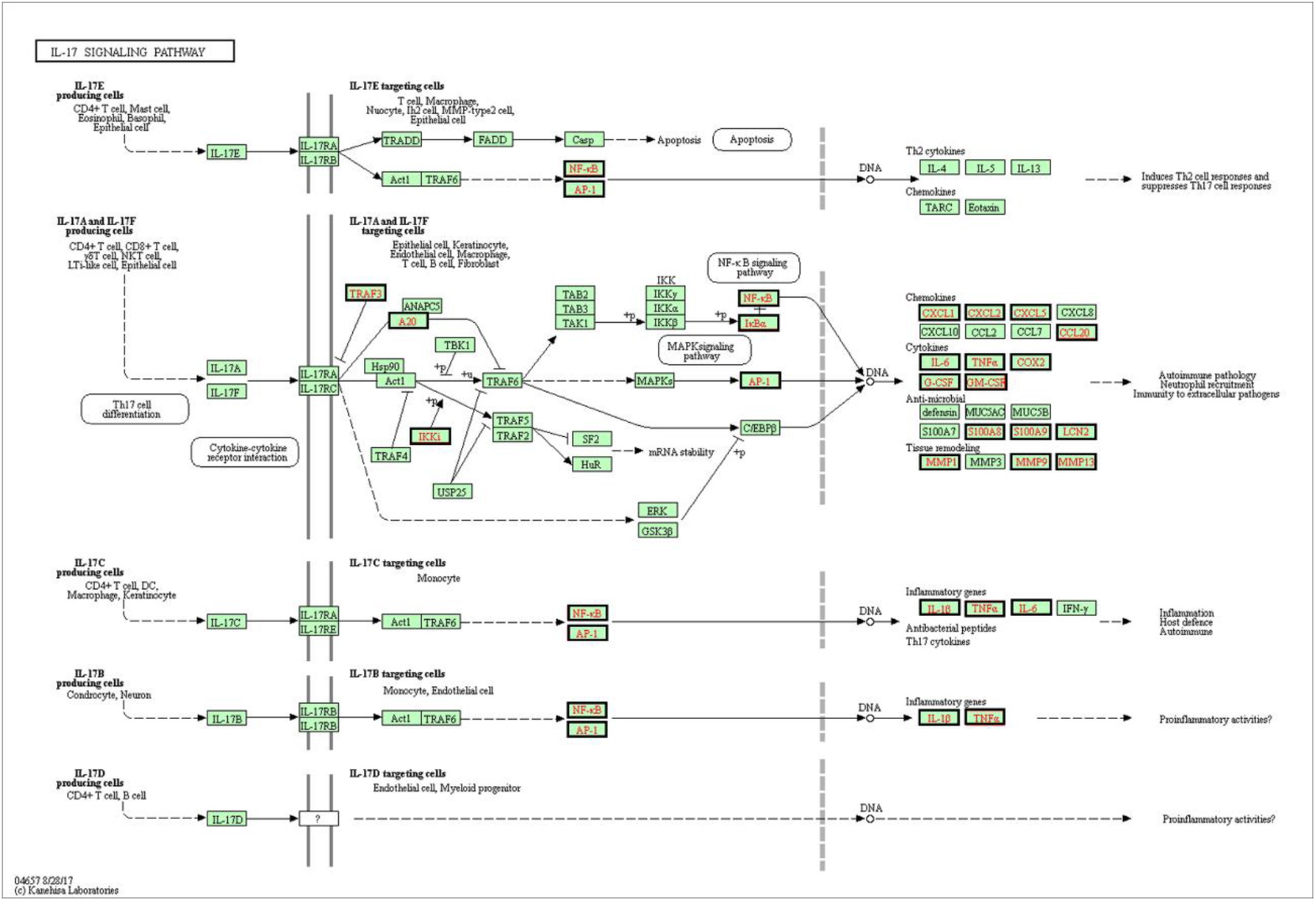
IL-17 signalling pathway. Genes significantly upregulated by SARS-CoV-2 infection are shown in bold frames.

**Figure 3.**
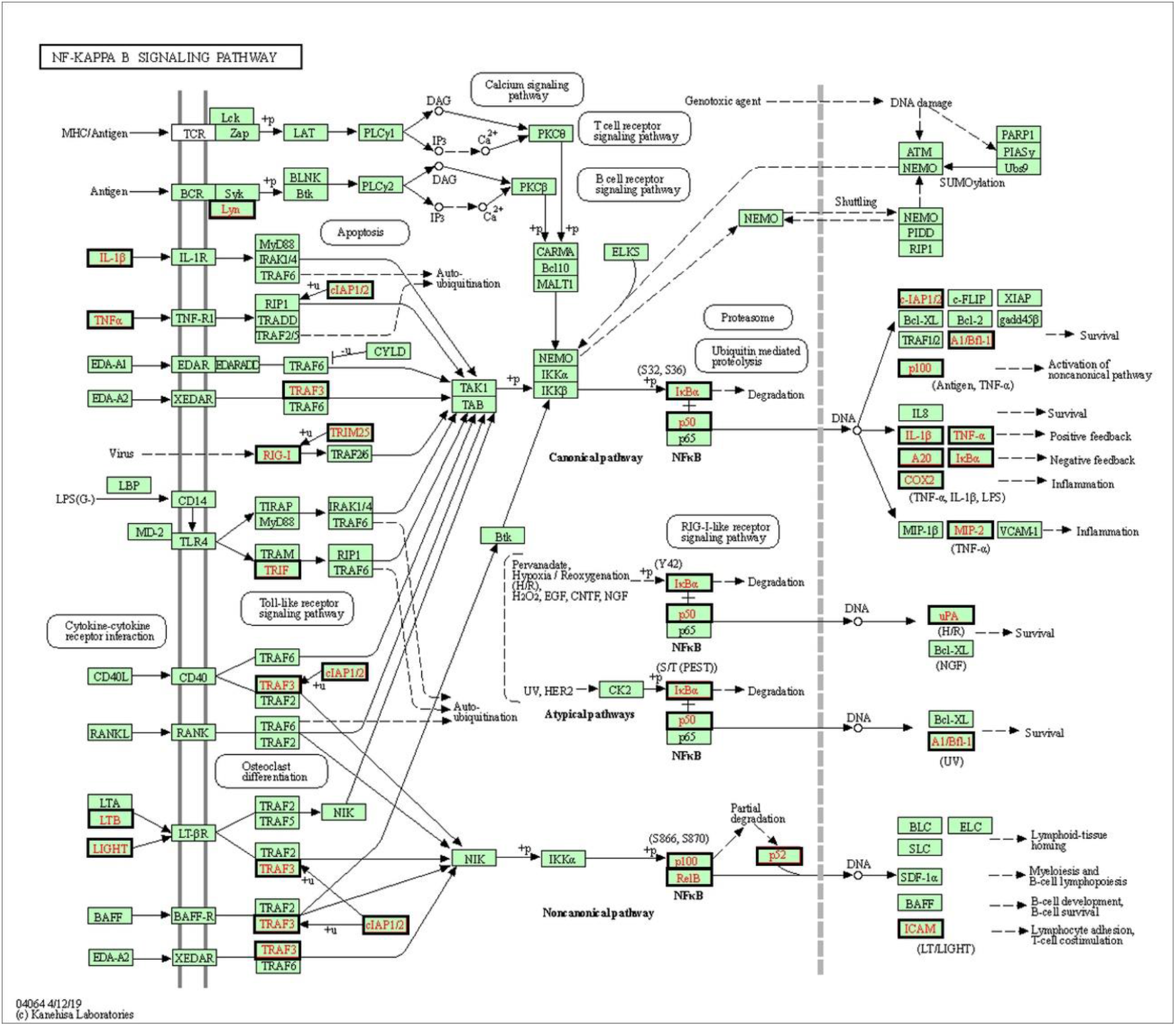
NFκB signalling pathway. Genes significantly upregulated by SARS-CoV-2 infection are shown in bold frames.

**Table 1.** GO biological processes enriched in significantly upregulated genes between SARS-CoV-2 positive and control NHBE cells.

| **GO biological processes** | **Ref** | **Hit** | **FDR** | **Query** |
| --- | --- | --- | --- | --- |
| **Immune system process** | 2769 | 142 | 1.36E-36 | 320 |
| **Defense response** | 1360 | 101 | 3.08E-36 | 320 |
| **Response to cytokine** | 1121 | 92 | 8.87E-36 | 320 |
| **Response to external biotic stimulus** | 1349 | 99 | 2.40E-35 | 320 |
| **Response to other organism** | 1347 | 99 | 2.66E-35 | 320 |
| **Response to biotic stimulus** | 1381 | 99 | 1.31E-34 | 320 |
| **Immune response** | 1870 | 113 | 5.61E-34 | 320 |
| **Response to external stimulus** | 2472 | 129 | 9.38E-34 | 320 |
| **Response to stress** | 3620 | 156 | 1.64E-33 | 320 |
| **Cellular response to cytokine stimulus** | 1031 | 83 | 1.30E-31 | 320 |
| **Cytokine-mediated signalling pathway** | 694 | 67 | 5.22E-29 | 320 |
| **Interspecies interaction between organisms** | 1997 | 108 | 2.71E-28 | 320 |
| **Response to organic substance** | 3025 | 133 | 8.18E-28 | 320 |
| **Defense response to other organism** | 967 | 75 | 2.95E-27 | 320 |
| **Innate immune response** | 776 | 66 | 1.13E-25 | 320 |
| **Response to stimulus** | 8544 | 226 | 9.05E-23 | 320 |
| **Cellular response to organic substance** | 2396 | 109 | 1.22E-22 | 320 |
| **Cell surface receptor signalling pathway** | 2480 | 111 | 1.25E-22 | 320 |
| **Cellular response to chemical stimulus** | 2956 | 122 | 1.64E-22 | 320 |
| **Response to chemical** | 4501 | 155 | 2.25E-22 | 320 |
| **Inflammatory response** | 505 | 49 | 1.06E-20 | 320 |
| **Regulation of response to external stimulus** | 1011 | 67 | 1.69E-20 | 320 |
| **Immune effector process** | 1079 | 67 | 4.71E-19 | 320 |
| **Regulation of response to stress** | 1452 | 77 | 1.88E-18 | 320 |
| **Response to virus** | 286 | 35 | 3.76E-17 | 320 |

**Table 2.** KEGG Pathways enriched in significantly upregulated genes between SARS-CoV-2 positive and control NHBE cells.

| **KEGG Pathways** | **Ref** | **Hit** | **FDR** | **Query** |
| --- | --- | --- | --- | --- |
| **IL-17 signalling pathway** | 92 | 25 | 1.32E-17 | 320 |
| **NF-kappa B signalling pathway** | 100 | 23 | 1.30E-14 | 320 |
| **Influenza A** | 167 | 28 | 2.24E-14 | 320 |
| **TNF signalling pathway** | 112 | 23 | 9.40E-14 | 320 |
| **Rheumatoid arthritis** | 88 | 17 | 1.26E-09 | 320 |

Contrary to infection-related biological functions of the upregulated genes, the biological functions of downregulated genes are related with organ development, especially kidney development as shown in Supplementary Table 1. Although adjusted P-value of enrichment in kidney development pathway is relatively high, and not many genes in this pathway are upregulated (9/272), this result may indicate a biologically relevant phenomenon, as a connection between SARS-CoV-2 and acute kidney injury is recently reported (Fanelli et al., 2020). Acute renal dysfunction has been observed as a comorbidity in 29% of the patients in a small cohort of 52 patients (Yang et al., 2020), a meta-analysis of data from 34 publications indicated that existing chronic kidney disease and acute kidney injury are strongly correlated with increased disease severity in Covid-19 patients (Wang et al., 2020). Further, secondary effects of the infection such as low levels of oxygen being delivered to organs or cytokine storm leading to clots may worsen the kidney dysfunction with adverse effects on the patient.

It has been also reported that respiratory viral infections increase the incidence of rheumatoid arthritis (RA). To date, three viruses which increases the RA incidence significantly were identified, one of which is SARS-CoV-2 (Joo et al., 2019). The genes upregulated in RA pathway response to SARS-CoV-2 are provided in Supplementary Figure 2.

Unsurprisingly, Influenza A signalling pathway was also found to be upregulated significantly. However, virus release pathway of Influenza A does not seem to be affected transcriptionally, this may mean that virus release follows an alternative route in SARS-CoV-2 infection in NHBE cells. Influenza A signalling pathway can be seen in Figure 4, the transcriptional response of the pathway in Covid-19 and swine flu is discussed in below sections. Other pathways that play important roles in initiating the inflammation are also discussed below.

**Figure 4.**
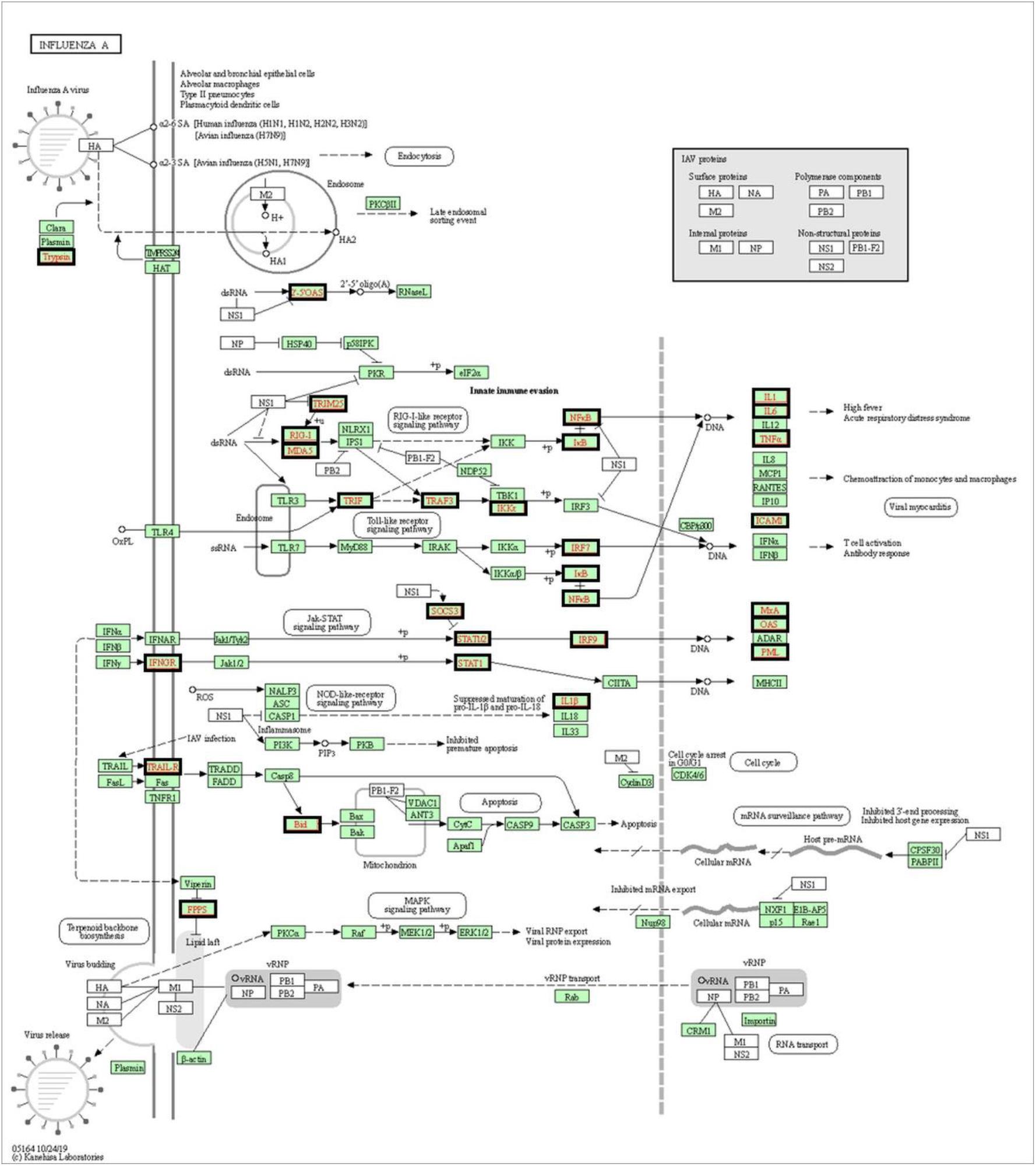
Influenza A signalling pathway. Genes significantly upregulated by SARS-CoV-2 infection are shown in bold frames.

### 3.1. IL-17 Signalling pathway

One of the most upregulated pathways in NHBE cells in response to SARS-CoV-2 infection is interleukin 17 (IL-17) signalling pathway, IL-17 is strongly associated with immunopathology but it also has important roles in host defence (Veldhoen, 2017).

Upregulated genes that encode the members of the IL-17 signalling pathway are shown in Figure 2, interleukins are usually cell type specific and produced by T cells as summarized in the figure. There are several subgroups of T helper cells such as Th1, Th2, Th9, Th17, T22 and T follicular helper (Tfh) cells (Linterman and Vinuesa 2010; Chen et al., 2019; Mousset et al., 2019). Th1 cells are responsible for activation of macrophages, cell-mediated immunity and also they increase the expression level of tolllike receptors (TLRs), which are part of the innate immune system. Th2 responsible for the production of antibodies and activation of eosinophils, supporting the host defence against extracellular organisms including helminths. Th17 cells secrete IL-17 themselves and play a critical role in the immune response to pathogens and several inflammatory diseases such as rheumatoid arthritis, asthma, allergy (Rahimi et al., 2019). The branch of IL-17 signalling pathway triggered by IL-17E which induces Th2 production and supress Th17 production is not activated transcriptionally, as expected. However, most targets of the branch triggered by IL-17A and IL-17F are upregulated. This branch of the signalling pathway induces Th17 cells, which in turn regulates autoimmune pathology, neutrophil recruitment and the immunity to extracellular pathogens. Neutrophils are also part of the innate immune system, which generate the early response to infections and activate adaptive immune system by antigen presentation.

Chemokines such as CXCL1, CXCL2, CXCL5 have chemotactic activity in neutrophils, they play role in inflammation and target injured or infected tissues (Li et al., 2019). Upregulated MMPs and chemokines demonstrate the response generated by NHBE cells to trigger the immune system, these two subgroups of genes are further discussed below comparatively in SARS-CoV-2 and H1N1 infection.

IL-1β and TNFα are also among the upregulated targets of IL-17 pathway, these two cytokines activate the canonical NFκB pathway. The transcription factor NFκB itself is activated by the IL-17 pathway and upregulates transcription of its targets, hence, forms a positive feedback loop in response to viral infection. Interestingly, none of the IL-17 ligands were expressed in detectable amounts in NHBE cells, and none of the ligands or receptors were upregulated in response to virus infection, therefore the positive feedback loop is likely to be initiated by activation of NFκB via other signalling pathways.

### 3.2. NF-Kappa B Signalling Pathway

NFκB signalling pathway is a proinflammatory signalling pathway with NFκB as its key transcription factor. NF-κB is a member of Rel-related protein family. This family also includes RelA (p65), RelB, c-Rel, NFκB1 (p105/p50) and NFκB2 (p100/p52). The target genes of NFκB are (TNF)-α, IL-2, IL6, IL-8 and interferon-β which play role in immune response and inflammation (Liao et al. 2005). Thus, it has critical role in pathogenesis and lung diseases. This pathway is composed of typically two branches, these are canonical, and non-canonical branches (Liao et al., 2005; Dosch et al., 2009). Canonical NFκB signalling pathway is implicated in pathogenesis of inflammatory bowel disease (IBD), asthma, chronic obstructive pulmonary disease (COPD) and RA (Lawrence, 2009). Unlike canonical NFκB, non-canonical NFκB is associated with development processes, such as B-cell survival and maturation of B-cell, bone metabolism and dendritic cell activation. In addition, the activation of canonical part of this pathway is faster but less persistent than non-canonical NFκB (Sun, 2017).

Upregulated genes in NFκB signalling pathway is shown in Figure 3, most targets of the canonical branch are upregulated, whereas only one target of the non-canonical branch is upregulated. Targets of the third branch, which is “atypical branch”, activated by a crosstalk of the two branches, were also upregulated in NHBE cells. It is plausible that infected tissues give priority to defence and survival rather than signalling for production of B cells. In this pathway, unlike the IL-17 signalling pathway, many ligands are upregulated although the receptors seem to be unaffected transcriptionally.

### 3.3. TNF Signalling Pathway

Tumour necrosis factor (TNF) plays important role in pro-inflammatory and anti-inflammatory processes. It provides protection against cancer and infectious pathogens. TNF signalling pathway includes 2 types of receptors, TNFR1 and TNFR2. The branch of signalling pathway which includes TNFR1 receptor lead to apoptosis, necroptosis and immune system related biological functions via crosstalk with other signalling pathways such as NFκB and MAPK. Its role is implicated in immune system homeostasis, antitumour responses, and control of inflammation (Mehta et al., 2018). Signalling via TNFR1, activates CD4 and CD8 T cells or innate immune system cells which induce the death of infected cells (Kalliolias and Ivashkiv, 2016; Mehta et al., 2018). TNFR2 expression is limited with some cell types such as neurons, immune cells and endothelial cells. TNFR2 promotes homeostatic effects locally such as cell survival and tissue regeneration (Kalliolias and Ivashkiv, 2016).

TNF, a target of NFκB signalling pathway is upregulated in infected NHBE cells, triggering the TNF signalling pathway (Figure 5). Most targets of the pathway such as inflammatory cytokines are transcriptionally upregulated including the cytokines involved in leukocyte recruitment.

**Figure 5.**
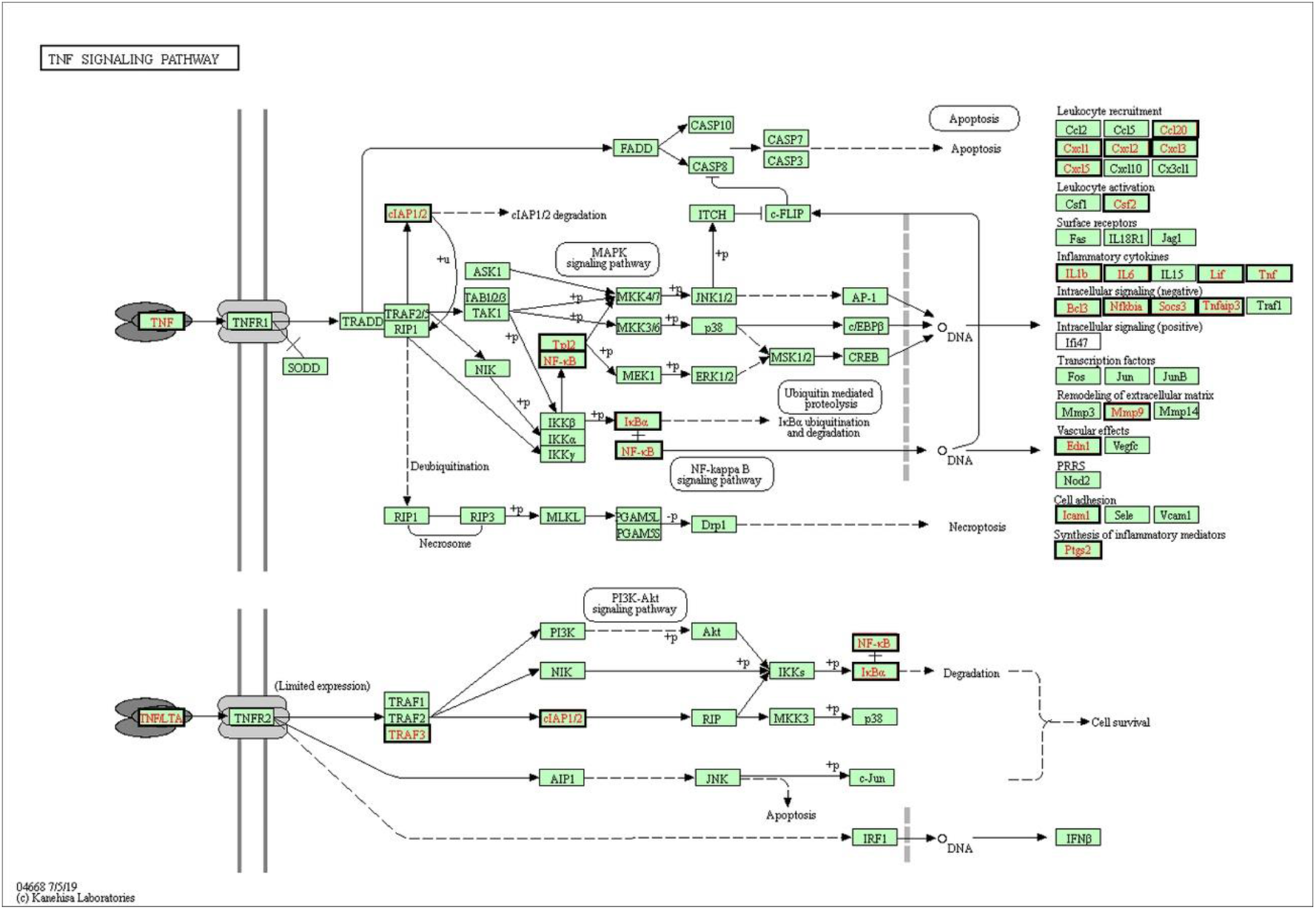
TNF signalling pathway. Genes significantly upregulated by SARS-CoV-2 infection are shown in bold frames.

### 3.4. ACE2 and TMPRSS2

Angiotensin converting enzyme 2 (ACE2) is expressed in human airway epithelia. Coronaviruses enter target cells via binding of their spike proteins to ACE2 proteins of the target cells. Therefore, the expression level of ACE2 has an impact on the possibility of infection transmission (Jia et al., 2005; Hoffmann et al., 2020). It is reported that overexpression of human ACE2 increases the severity of SARS-CoV in infected mice, and the expression of ACE2 decreases after infection. ACE2, which is then re-injected into mice, causes high levels of lung injury (Kuba et al., 2005). Spike protein of SARS-CoV-2 and SARS-CoV share ~76% amino acid identity, and expression level of ACE2 may also have an impact on SARS-CoV-2 infection (Hoffmann et al., 2020). According to our differential gene expression analysis results, ACE2 did not change its expression significantly. A cellular protease, TMPRSS2, reported to take part in mediating the virus entry, also had invariant expression levels in NHBE cells. NHBE cells produced a strong transcriptional response to SARS-CoV-2 although counts from the RNA-Seq data analysed here indicated very low expression of ACE2 gene in the cells, as previously reported by Blanco-Melo et al. (2020). It is therefore not clear whether if very small amounts of ACE2 protein is enough to mediate the infection or another unknown protein also takes part. Further, our results imply that ACE2 or TMPRSS2 transcription is not regulated by the virus infection either negatively or positively in bronchial epithelial cells.

### 3.5. Comparison of SARS-CoV-2 and H1N1 infection in NHBE cells

H1N1 infection which causes swine flu (Influenza A) is lethal only in 0.1 – 0.02% of the patients, as opposed to anticipated 1-3% lethality in Covid-19. We analysed the transcriptional response in NHBE cells to H1N1 infection, then compared the response to that of SARS-CoV-2.

Gene expression levels from four replicates of H1N1 infected and control cells were compared, 1899 genes were found to be upregulated and 474 genes were found to be downregulated in NHBE cells 12 hours after the H1N1 infection (Padj < 0.05). GO terms and KEGG pathways that were enriched with upregulated genes were similar to those of SARS-CoV-2, but we found additional terms such as metabolism and ribosomes, indicating that impact of H1N1 infection was not limited to virus response (Supplementary Table 2 and 3). We mapped the upregulated and downregulated genes on KEGG Influenza A pathway (Supplementary Figure 3), only 4 genes were downregulated, while about a quarter (41/167) of the genes that have roles in the pathway or that are targeted by the pathway were upregulated, as expected.

We compared the transcriptional response to SARS-CoV-2 and H1N1 infection in NHBE cells to elucidate the differential gene expression in Covid-19 in comparison to swine flu, 78 genes were upregulated by both virus infections, whereas only 3 genes were downregulated by both. 17 of the 78 upregulated genes were members of the Influenza A pathway, indicating a limited shared response on the pathway.

Next, we focused on SARS-CoV-2-specific transcriptional response. Out of 24 genes upregulated in SARS-Cov-2 infection in the IL-17 pathway, only 6 was also upregulated in H1N1 infection. Similar comparison for other pathways has shown that 9 out of 23 in NFκB, 9 out of 23 in TNF, 4 out of 17 in RA and 17 out of 28 in Influenza A pathways were shared in both infections. Virus response pathways had a stronger transcriptional response to SARS-CoV-2 infection than to H1N1 infection, except for Influenza A pathway, which experienced a stronger response to H1N1 infection, as expected (Supplementary Table 3). Although some of the differences observed between the responses may be attributed to sampling being done after 24 hours in SARS-CoV-2 as opposed to sampling being done after 12 hours in H1N1 infection, the expected response profile in Influenza A pathway suggest that sampling time has a minor effect on the results.

### 3.6. Comparison of cytokine and MMP expression in SARS-CoV-2 and H1N1 infection

Finally, we compared expression levels of a subset of genes which encode the protein that are secreted to extracellular matrix in SARS-CoV-2 and H1N1 infection (Figure 6). Cytokine CSF3 was very significantly upregulated in response to SARS-CoV-2 infection, but was invariant in response to H1N1 infection. Also, IL36A, CXCL5 and CSF2 had high fold changes in SARS-CoV-2 infection when compared to H1N1 infection (Figure 6A, B).

**Figure 6.**
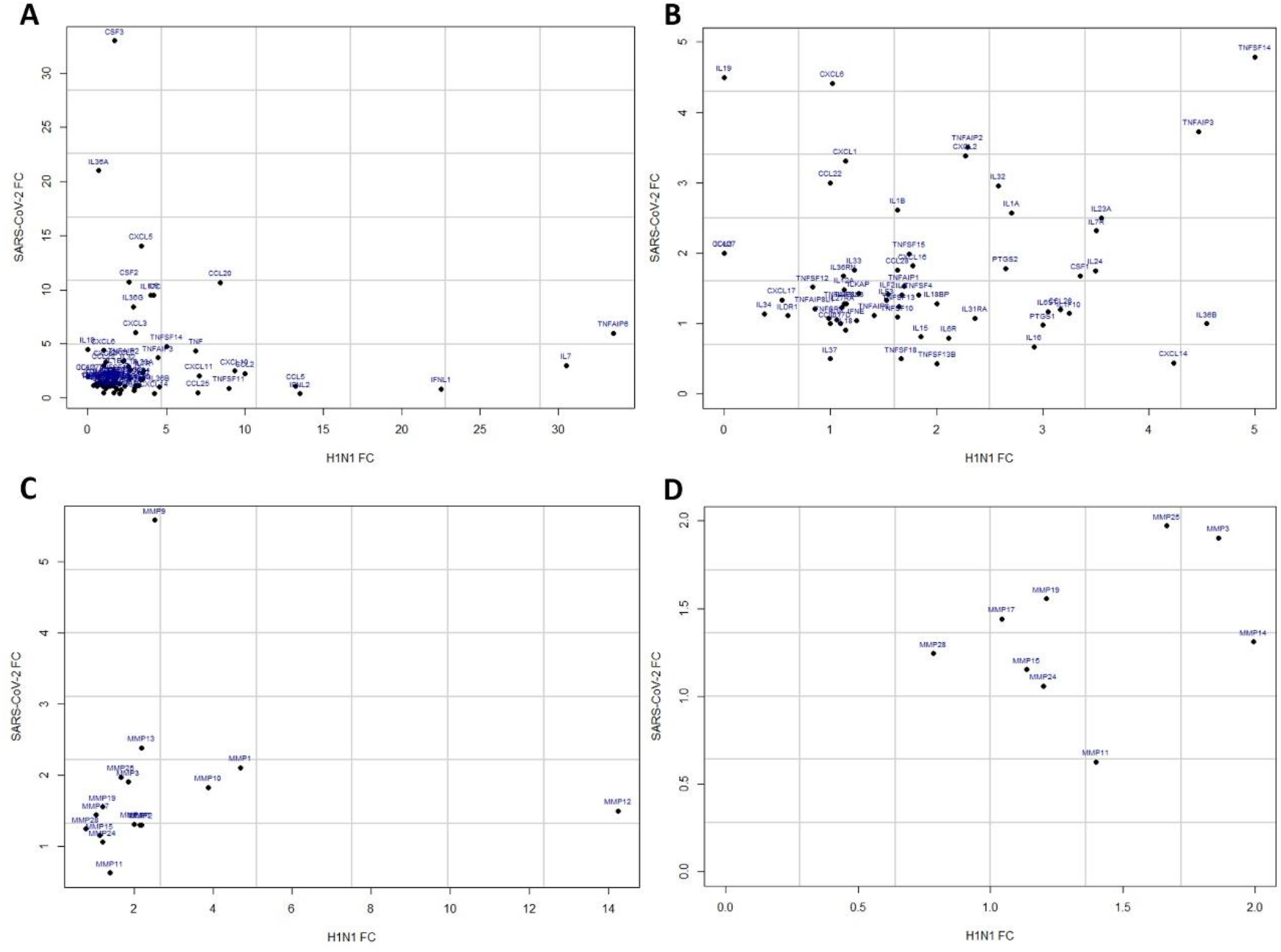
Fold changes of cytokines (A and B) and matrix metal proteases (C and D) in SARS-CoV-2 (y-axis) and H1N1 (x-axis) infected cells when compared to control cells. B and D provides a zoom into A and C respectively, B is [0-5, 0-5] range of A and D is [0-2, 0-2] range of C.

Blood samples collected from 41 patients in acute phase of Covid-19 were analysed for levels of 27 cytokines (Wang et al. 2020), CSF3 was one of three cytokines which had significantly different expression in healthy individuals, patients in intensive care and patients not in intensive care, hence CSF3 had a discriminative power not only between healthy and disease cases, but also between severe and less severe disease cases. The other two cytokines were TNFα and MIP1α, former had similar fold changes in SARS-CoV-2 and H1N1 infections, later was not included in our dataset.

The samples were collected 24 hours after the SARS-CoV-2 infection to generate the dataset analysed in this study, therefore the data reflects the early response to infection. 30 fold change in expression of CSF3 in NHBE cells in 24 hours may mean that this cytokine is an early marker of the infection and potentially can be detected in blood samples before the patients develop any symptoms of the disease.

Another subgroup we analysed are matrix metalloproteinases (MMPs), proteolytic enzymes which play multiple roles in immune response, tissue degradation and regulation of inflammation via activation and inactivation of cytokines and chemokines (Elkington et al., 2005; Dandachi and Shapiro, 2014). Transcriptional response of MMP genes in SARS-CoV-2 versus H1N1 infections are shown in Figure 6C and D, MMP9 and MMP12 responded differentially in two types of infection. MMP9 is involved in respiratory epithelial healing and its level is elevated in rheumatoid arthritis, whereas MMP12 may take part in development of emphysema, an obstructive lung disease.

Both CSF3 and MMP9 are targets of IL-17 pathway, MMP9 is also a target of TNF pathway, hence upregulation of these two genes are likely to be a result of activities of IL-17 and TNF pathways.

### 3.7. Crosstalk of signalling pathways

Our analyses demonstrated that multiple immune response pathways are activated in response to SARS-CoV-2 infection in NHBE cells. In the *in vitro* system we analysed, no external signals are available and NHBE cells do not express ligands of IL-17 pathway themselves, although targets of the pathway are significantly upregulated. Two of those, TNFα and IL1β, mediate the positive feedback loops with TNFα and NFκB pathways. In Figure 7, targets of the three pathways are shown, only the upregulated proteins targeted by at least two pathways are shown for clarity. The crosstalk between these pathways are likely to produce a robust response to infection via positive feedback loops, where TNFα and IL1β play a central role together with the NFκB pathway. Activity of these pathways lead to high expression of cytokines such as CSF3 and metalloproteases such as MMP9 which in turn would orchestrate the recruitment of immune cells such as leukocytes in *in vivo* tissues. ICAM1, located on membranes of leukocytes and endothelial cells is mostly expressed by endothelial cells. Indeed, in NHBE cells, we find that TNF and NF-κB signalling pathways co-activate ICAM1 expression in response to SARS-CoV-2 infection. Under in vivo conditions. ICAM1 would mediate attachment of leukocytes to the endothelial cells as part of the immune response (Frank and Lisanti, 2008). Similarly, regulation by the three pathways converge in activation of inflammatory response related cytokines such as CXCL1, CXCL2, CXCL3 and PTGS2.

**Figure 7.**
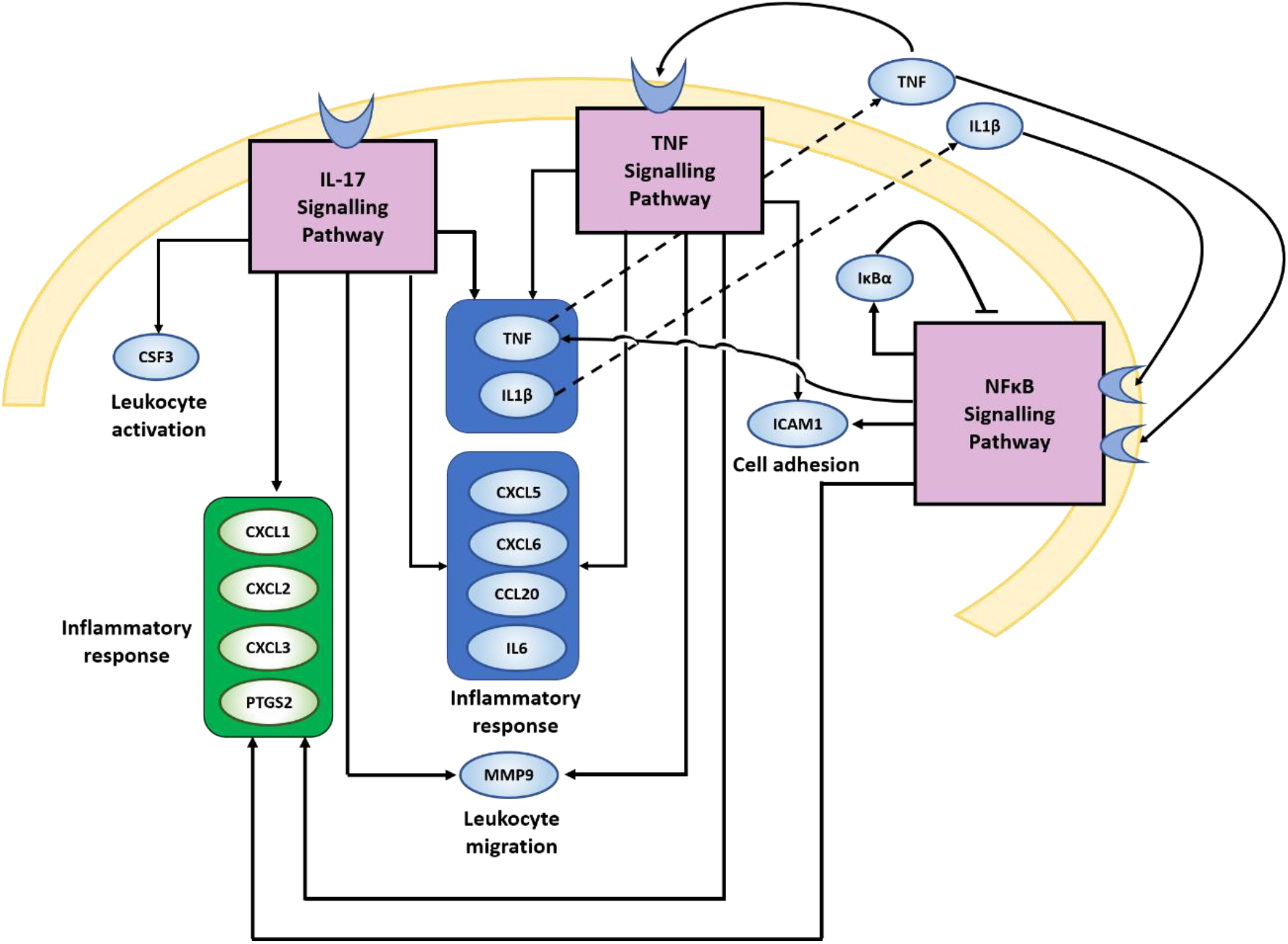
Crosstalk of signalling pathways in SARS-CoV-2 infected NHBE cells.

## 4. DISCUSSION

There is an ongoing global effort to better understand the Covid-19 disease caused by SARS-CoV-2 to be able to develop high sensitivity / high specificity diagnosis tests and better treatment methods. In this study, we examined the signalling pathways triggered by SARS-CoV-2 infection and found that NFκB is central to upregulated signalling pathways. Our study is limited to response generated in 24 hours in NHBE cells grown in vitro. However, analysis of the early response of the cells gave important clues about the cytokine storm experienced by most patients which increases the severity of the disease. We identified a subgroup of genes related to renal system disorders and rheumatoid arthritis which also respond to SARS-CoV-2 infection.

Further, we compared the transcriptional response to SARS-CoV-2 and H1N1 infection in NHBE cells. We find that response generated in H1N1 infection is much stronger in the Influenza A pathway as expected, but other immune response pathways such as IL-17, NFκB and TNF pathways yielded a stronger response in SARS-CoV-2 infection, which may explain the severity and higher mortality rates of inflammation in Covid-19 patients when compared to swine flu. We identified a cytokine, CSF3, and a matrix metal protease, MMP9 to be exclusive to SARS-CoV-2 when compared to H1N1. These two extracellular proteins should be further investigated as potential markers of the infection in blood samples.

This study is limited with samples collected at two time points and infection by two viruses in NHBE cells. Time course data from other types of infections and data from patient samples are needed to confirm the results we present here. We believe a thorough mechanistic understanding of virus response and signalling crosstalk will lead to new possibilities in developing test methods and treatment options for Covid-19.

## Acknowledgements

Enes Ak is funded by The Scientific and Technological Research Council of Turkey (TÜBİTAK) 116S388 and The Turkish Directorate of Health Institutes (TÜSEB) 2019-TA-01-3936. We thank O.Veli for the list of cytokines.

---

**Supplementary Figure 1.**
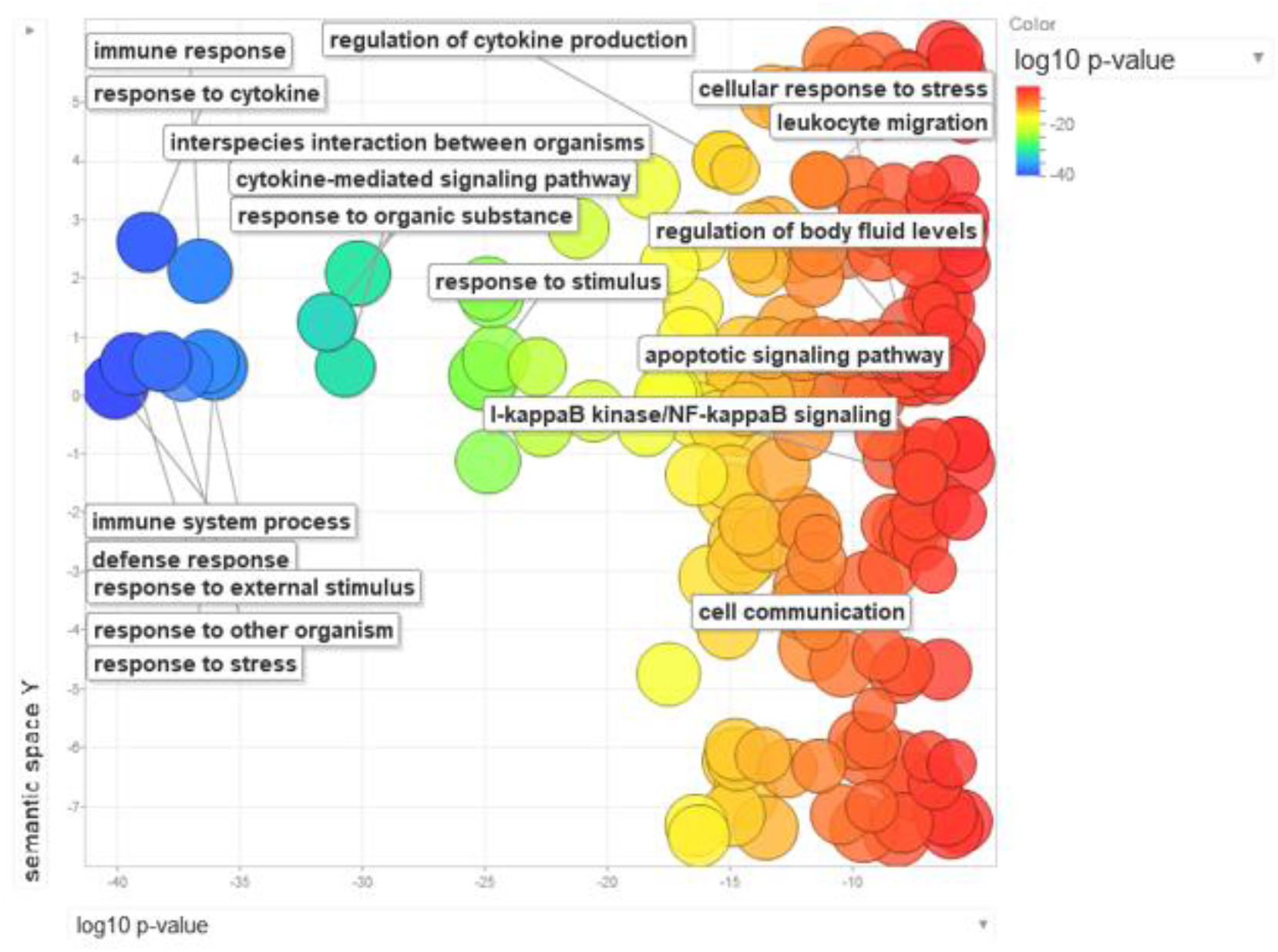
The visualization of GO biological processes results of upregulated significantly changed genes between SARS-CoV-2 positive and control NHBE cells. The x-axis represents log10(p-value) and the y-axis represents semantic relatedness between GO terms. Therefore, while left side of the figure gives most significant GO terms, right side gives less significant results. Proximity of two terms on the y-axis indicate that they are biologically related to each other.

**Supplementary Figure 2.**
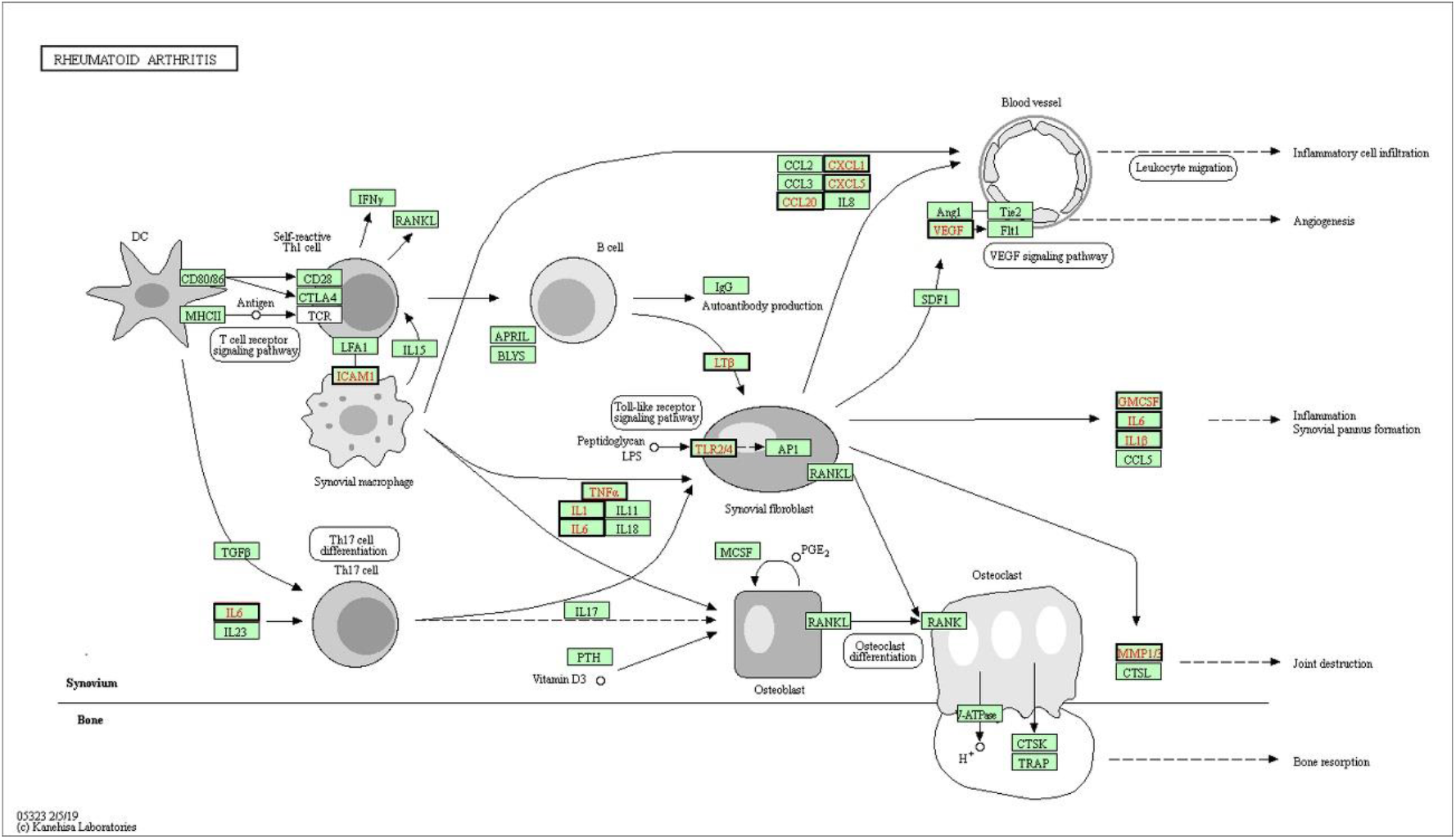
Rheumatoid arthritis signalling pathway. Genes significantly upregulated by SARS-CoV-2 infection are shown in bold frames.

**Supplementary Figure 3.**
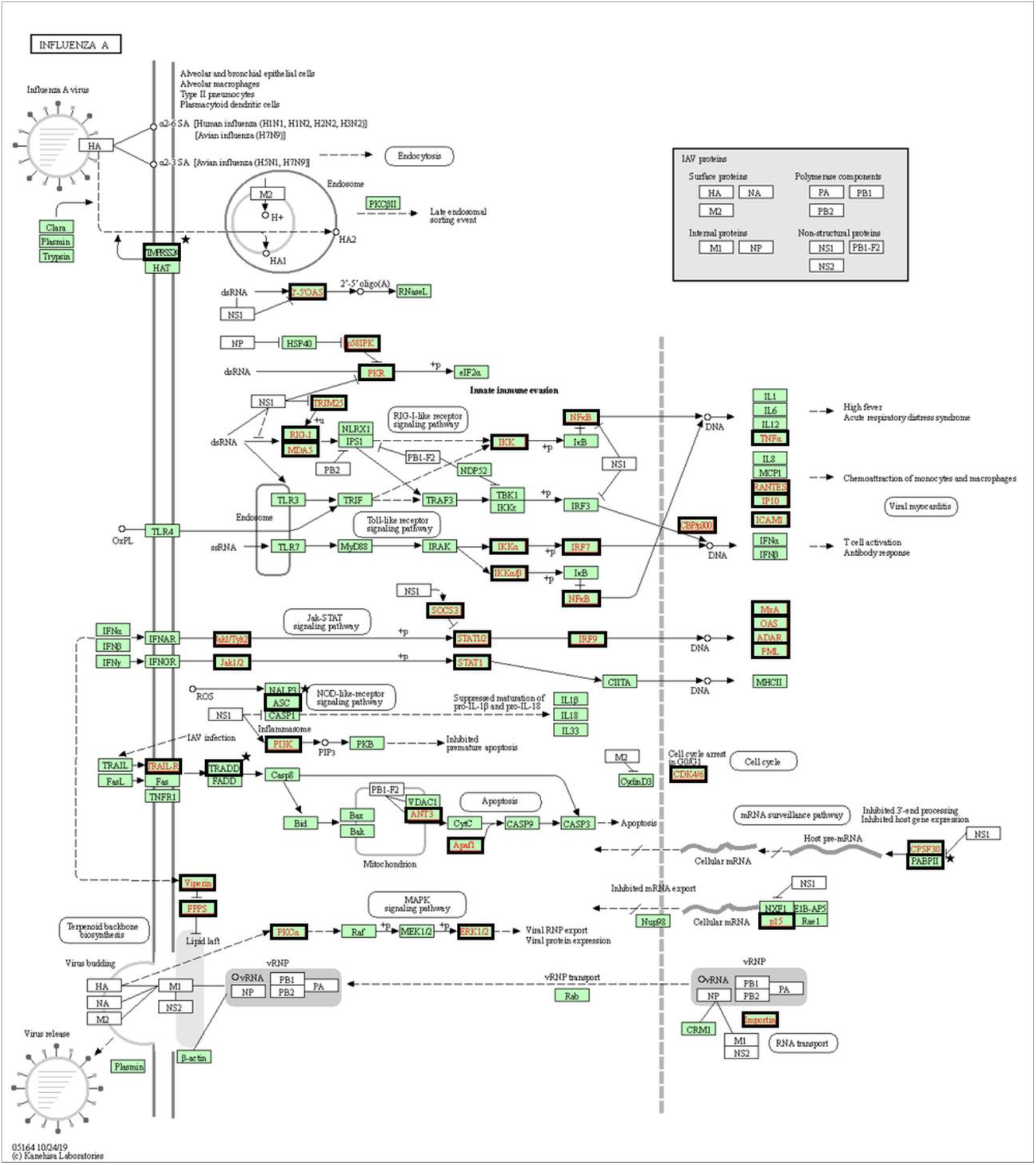
Influenza A signalling pathway. Genes significantly upregulated are shown in bold frames and downregulated by H1N1 infection are shown in bold frames and with stars.

**Supplementary Table 1.** GO biological processes enriched in significantly downregulated genes between SARS-CoV-2 positive and control NHBE cells.

| **GO biological processes** | **Ref** | **Hit** | **FDR** | **Query** |
| --- | --- | --- | --- | --- |
| **Animal organ development** | 3215 | 41 | 4.03E-03 | 115 |
| **Multicellular organism development** | 5065 | 52 | 2.14E-02 | 115 |
| **System development** | 4444 | 47 | 2.38E-02 | 115 |
| **Developmental process** | 5900 | 57 | 2.56E-02 | 115 |
| **Positive regulation of biological process** | 6249 | 59 | 2.71E-02 | 115 |
| **Anatomical structure development** | 5451 | 53 | 2.86E-02 | 115 |
| **Platelet degranulation** | 128 | 7 | 3.03E-02 | 115 |
| **Regulation of developmental process** | 2631 | 32 | 3.40E-02 | 115 |
| **Biological process** | 17794 | 115 | 3.54E-02 | 115 |
| **Unclassified** | 3057 | 3 | 3.99E-02 | 115 |
| **Multicellular organismal process** | 7015 | 62 | 4.35E-02 | 115 |
| **Kidney development** | 272 | 9 | 4.44E-02 | 115 |
| **Renal system development** | 282 | 9 | 4.93E-02 | 115 |

**Supplementary Table 2.**
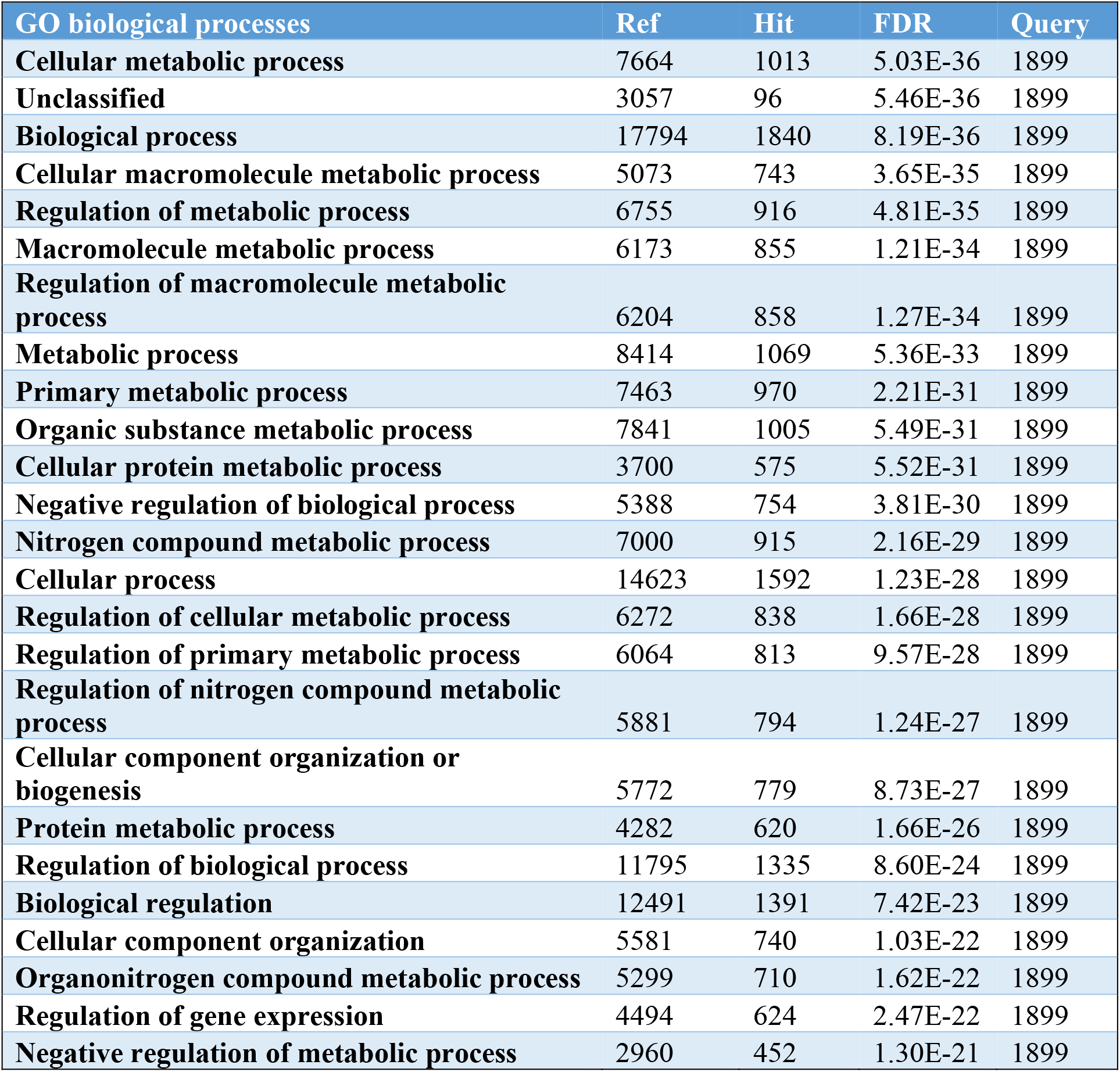
GO biological processes enriched in significantly upregulated genes between H1N1 positive and control NHBE cells.

**Supplementary Table 3.** KEGG Pathways enriched in significantly upregulated genes between H1N1 positive and control NHBE cells.

| KEGG Pathways | Ref | Hit | FDR | Query |
| --- | --- | --- | --- | --- |
| Ribosome | 134 | 48 | 1.44E-11 | 1899 |
| Protein processing in endoplasmic reticulum | 166 | 45 | 1.61E-06 | 1899 |
| Hepatitis C | 155 | 41 | 1.03E-05 | 1899 |
| Human papillomavirus infection | 330 | 68 | 2.95E-05 | 1899 |
| Influenza A | 167 | 41 | 5.44E-05 | 1899 |
| Proteoglycans in cancer | 202 | 45 | 2.34E-04 | 1899 |
| Pancreatic cancer | 76 | 23 | 2.61E-04 | 1899 |
| mTOR signaling pathway | 152 | 35 | 9.13E-04 | 1899 |
| Alzheimer disease | 171 | 38 | 9.13E-04 | 1899 |
| Kaposi sarcoma-associated herpes virus infection | 186 | 40 | 1.06E-03 | 1899 |
| Viral carcinogenesis | 199 | 42 | 1.06E-03 | 1899 |
| Hepatitis B | 162 | 36 | 1.11E-03 | 1899 |
| Measles | 138 | 32 | 1.13E-03 | 1899 |
| Human cytomegalovirus infection | 223 | 45 | 1.41E-03 | 1899 |
| Focal adhesion | 198 | 41 | 1.51E-03 | 1899 |
| TNF signaling pathway | 112 | 27 | 1.76E-03 | 1899 |
| Renal cell carcinoma | 68 | 19 | 2.31E-03 | 1899 |
| Prostate cancer | 97 | 24 | 2.47E-03 | 1899 |
| Non-alcoholic fatty liver disease (NAFLD) | 149 | 32 | 3.33E-03 | 1899 |
| Pathways in cancer | 529 | 86 | 3.33E-03 | 1899 |
| Chronic myeloid leukemia | 76 | 20 | 3.33E-03 | 1899 |
| NOD-like receptor signaling pathway | 177 | 36 | 4.06E-03 | 1899 |
| EGFR tyrosine kinase inhibitor resistance | 79 | 20 | 4.92E-03 | 1899 |
| Colorectal cancer | 86 | 21 | 5.76E-03 | 1899 |
| TGF-beta signaling pathway | 93 | 22 | 6.29E-03 | 1899 |

**Supplementary Table 4.** GO biological processes enriched in significantly downregulated genes between H1N1 positive and control NHBE cells.

| GO biological processes | Ref | Hit | FDR | Query |
| --- | --- | --- | --- | --- |
| Organic substance biosynthetic process | 2886 | 110 | 1.02E-03 | 474 |
| Cellular biosynthetic process | 2782 | 106 | 1.30E-03 | 474 |
| Biosynthetic process | 2964 | 113 | 1.32E-03 | 474 |
| G protein-coupled receptor signaling pathway | 1322 | 7 | 1.85E-03 | 474 |
| Mitochondrial translational elongation | 88 | 12 | 8.35E-03 | 474 |

